# Poly-alanine-tailing is a modifier of neurodegeneration caused by Listerin mutation

**DOI:** 10.1101/2024.08.24.608776

**Authors:** Hao-Chih Hung, Carlos Costas-Insua, Sarah E. Holbrook, Jennifer E. Stauffer, Paige B. Martin, Tina A. Müller, David G. Schroeder, Yu Kigoshi-Tansho, Haifei Xu, Rüdiger Rudolf, Gregory A. Cox, Claudio A. P. Joazeiro

## Abstract

The surveillance of translation is critical for the fitness of organisms from bacteria to humans. Ribosome-associated Quality Control (RQC) is a surveillance mechanism that promotes the elimination of truncated polypeptides, byproducts of ribosome stalling during translation. In canonical mammalian RQC, NEMF binds to the large ribosomal subunit and recruits the E3 ubiquitin ligase Listerin, which marks the nascent-chains for proteasomal degradation. NEMF additionally extends the nascent-chain’s C-terminus with poly-alanine (‘Ala-tail’), exposing lysines in the ribosomal exit tunnel for ubiquitination. In an alternative, Listerin-independent RQC pathway, released nascent-chains are targeted by Ala-tail-binding E3 ligases. While mutations in Listerin or in NEMF selectively elicit neurodegeneration in mice and humans, the physiological significance of Ala-tailing and its role in disease have remained unknown. Here, we report the analysis of mice in which NEMF’s Ala-tailing activity was selectively impaired. Whereas the *Nemf* homozygous mutation did not affect lifespan and only led to mild motor defects, genetic interaction analyses uncovered its synthetic lethal phenotype when combined with the *lister* neurodegeneration-causing mutation. Conversely, the *lister* phenotype was markedly improved when Ala-tailing capacity was partially reduced by a heterozygous *Nemf* mutation. Providing a plausible mechanism for this striking switch from early neuroprotection to subsequent neurotoxicity, we found that RQC substrates that evade degradation form amyloid-like aggregates in an Ala-tail dependent fashion. These findings uncover a critical role for Ala-tailing in mammalian proteostasis, and deepen our molecular understanding of pathophysiological roles of RQC in neurodegeneration.

## Introduction

Protein translation is a fundamental cellular process stringently regulated and quality controlled (Schuller & Green, 2018; Joazeiro, 2019; Weisser & Ban, 2019; Korostelev, 2022). Faulty translation elongation causing ribosomes to stall can occur for various reasons, including translation into the polyA tail in transcripts lacking stop codons, or depletion of aminoacylated-tRNAs during starvation, among others (Moore & Sauer, 2007; Simms *et al*, 2014; Joazeiro, 2017; Chandrasekaran *et al*, 2019; Thomas *et al*, 2020; D’Orazio & Green, 2021; Stoneley *et al*, 2022; Yang *et al*, 2023; Suryo Rahmanto *et al*, 2023; Zhao *et al*, 2023; Vind *et al*, 2023; Snieckute *et al*, 2023; Sinha *et al*, 2024; Parker *et al*, 2024). Ribosome stalling poses a cellular threat because it not only sequesters ribosomes from the translationally-competent pool, but also produces partially-synthesized polypeptides that can be toxic (Defenouillere & Fromont-Racine, 2017; Joazeiro, 2019; Sitron & Brandman, 2020; Howard & Frost, 2021; Vind *et al*., 2023).

Mammalian cells detect, then rescue stalled ribosomes by splitting their subunits. This rescue reaction releases a large (60S) ribosomal subunit still obstructed with a nascent chain-tRNA conjugate (Shao *et al*, 2013; Juszkiewicz *et al*, 2018; Hashimoto *et al*, 2020; Juszkiewicz *et al*, 2020; Matsuo *et al*, 2023). These nascent chains are targeted for degradation by ribosome-associated quality control (RQC) (Bengtson & Joazeiro, 2010; Brandman *et al*, 2012; Defenouillere *et al*, 2013; Lyumkis *et al*, 2014; Shao *et al*, 2015). In RQC, the Nuclear Export Mediator Factor (NEMF, Rqc2 in yeast) first recognizes the 60S subunit obstruction and stabilizes the binding of the RING-domain E3 ligase Listerin, which ubiquitinates the nascent chain for proteasomal disposal (Sitron & Brandman, 2020; Filbeck *et al*, 2022; Inada & Beckmann, 2024).

Mammalian NEMF can further assist Listerin by elongating the nascent chain with an untemplated poly-alanine stretch (‘Ala-tail’), which helps expose lysine residues otherwise buried in the ribosome exit tunnel for ubiquitination (Shen *et al*, 2015; Kostova *et al*, 2017; Udagawa *et al*, 2021; Thrun *et al*, 2021). NEMF orthologs mediate C-terminal tailing by directly recruiting tRNA to the ribosomal A-site and promoting peptidyl-transfer, in a reaction mimicking canonical translation elongation (Shen *et al*., 2015; Osuna *et al*, 2017; Crowe-McAuliffe *et al*, 2021; Takada *et al*, 2021; Filbeck *et al*, 2021). In bacteria, the universally conserved D97 residue in the NEMF ortholog, RqcH (D96 in human NEMF) directly contacts the tRNA-Ala anticodon loop, and its mutation impairs tRNA-Ala binding and Ala-tailing, while sparing ribosome binding (Lytvynenko *et al*, 2019; Filbeck *et al*., 2021; Takada *et al*., 2021). Similarly, mutating the equivalent residue in Rqc2, D98, impairs C-terminal tailing in yeast (Shen *et al*., 2015; Yonashiro *et al*, 2016; Choe *et al*, 2016; Osuna *et al*., 2017).

We have recently uncovered that NEMF-mediated Ala-tailing also initiates a Listerin-independent degradation route (Thrun *et al*., 2021), whereby Ala-tails act as a *C*-terminal degron recognized by the E3 ligases Pirh2 and KLHDC10 (Thrun *et al*., 2021; Patil *et al*, 2023; Wang *et al*, 2023)., reminiscent of the RQC pathway in bacteria (Lytvynenko *et al*., 2019; Thrun *et al*., 2021). We named this novel RQC branch RQC-C after the *C*-terminal degron role of Ala tails (Thrun *et al*., 2021).

The (patho)physiological relevance of RQC is underscored by our earlier identification of a Listerin-mutant mouse line from a genome-wide N-ethyl-N-nitrosourea (ENU) mutagenesis screen (C57BL/6J *Ltn1* ^*liste*r*/lister*^, hereafter *Ltn1* ^*liste*r*/lister*^). These mice carry a splice donor site mutation in the *Ltn1* gene, resulting in in-frame deletion of a short exon and a loss-of-function Listerin variant missing 14 amino acids (Chu *et al*, 2009). The animals are born normal but develop severe, age-progressive motor neuron degeneration, muscle atrophy and motor dysfunction, and die prematurely. More recently, we found two additional ENU-derived mouse lines carrying homozygous mutations in *Nemf* (Martin et al, 2020), and displaying similar phenotypes to *lister*. Finally, we and others have found bi-allelic inactivating mutations in human NEMF strongly associated with neuromuscular disease in humans (Martin *et al*., 2020; Ahmed *et al*, 2021).

The above findings clearly implicate defective RQC activity in neurodegeneration. However, the physiological relevance of NEMF-mediated Ala-tailing during translational surveillance, and its role in neurodegeneration, have remained unknown. In this study, we investigate the role of mammalian Ala-tailing by developing a novel mouse model with a D96-to-A (D96A) mutation, designed to selectively impair NEMF’s Ala-tailing activity. Genetic interaction experiments demonstrate that Ala-tailing plays a protective role in translational homeostasis. Additionally, the data reveals that Ala-tailing contributes to neurotoxicity in *lister* mice. Cellular studies indicate that the accumulation of Ala-tailed proteins leads to the formation of SDS-resistant, amyloid-like aggregates, suggesting a molecular pathogenic mechanism. These findings expand our understanding of mammalian RQC and proteostasis mechanisms, and elucidate a novel molecular mechanism underlying neurodegeneration. Overall, this study highlights the critical roles of Listerin and NEMF activities in maintaining neuronal homeostasis, and the critical importance of this conserved protein quality control pathway.

## Materials and methods

### Generation of the Nemf-D96A mouse line

The *Nemf*-D96A strain was produced by CRISPR–Cas9 mutagenesis. Briefly, C57BL/6J zygotes were microinjected of with 100 ng/µL *Cas9* mRNA and 50 ng/µL of sgRNA targeting exon 4 of *Nemf*: 5’-GCTTGGTGTGGACAGAATTG-3’, and 20 ng/µL donor oligo *Nemf*: 5’-TTCTTGCTAGTGTCGAAAACATTTGAAGAGTCGGAGATTAGTCAGTGCAAAACAGCTTG GTGTGG**<u>C</u>**CAGAATTGTGGATTTCCAGTTTGGAAGTGACGAAGCTGCTTATCACTTAATC ATTGAGCTCTAT-3’ creating the D96A mutation GAC>GCC. Mosaic founder mice identified as carrying the targeted mutation were backcrossed to C57BL/6J and the progeny carrying the mutation was further backcrossed to C57BL/6J to establish the colony.

### Animals

All mouse husbandry and procedures were reviewed and approved by the respective Institutional Animal Care and Use Committees at The Jackson Laboratory and UF Scripps, and were carried out according to the NIH Guide for Care and Use of Laboratory Animals. Mice were bred and maintained under standard conditions (22 ºC, 42% humidity and a 12/12 dark/light cycle) with *ad libitum* access to food and water. For genotyping, DNA was extracted from an ear tissue punch using the Qiagen DNeasy blood and tissue kit (Cat# 69506). For *Nemf-D96A* allele genotyping, DNA was used as template in PCR with the following forward and reverse primers *NEMF*.*F: CATGGTGAATGGAGAGAACC*; *NEMF*.R: TTGATCCCAGCACTAGGGAG. Purified PCR products were analyzed by Sanger sequencing. The *lister* allele was genotyped as previously described (Chu *et al*., 2009). For genetic interaction experiments, animals were genotyped after weaning. For the intercross between *Ltn1* ^*lister*/*wt*^; *Nemf* ^D96A/D96A^ littermates, we monitored animals twice on the day of birth, and cadavers from animals dying at post-natal day 1 (*d 1*) were recovered and genotyped.

### Phenotypic and behavioral analyses

Mice were weighted weekly, starting at the second week post-birth. To measure muscle strength, we used the wire hang assay. Briefly, mice of indicated ages were placed on top of a wire mesh cage, which was immediately inverted, and the latency to fall was measured. Mice were removed after 60 s if they did not fall, and allowed to rest for a minimum of 5 minutes before repeating the test. Each value represents the maximum value obtained from three tests per timepoint. To assess the hind limb extension reflex, mice were suspended from the tail and the clenching response during the first 30 s was evaluated.

### Analysis of myelinated axons

The motor branch of the femoral nerve was dissected and fixed overnight at 4 °C in 2% glutaraldehyde and 2% paraformaldehyde in 0.1 M cacodylate buffer, followed by post-fixation in 1% osmium tetroxide in 0.1 M cacodylate buffer and embedding in Embed 812 Resin (Electron Microscopy Sciences, Hatfield, PA). One-µm wide sections were sliced on a Leica RM2265 rotary microtome equipped with a diamond knife. Tissue slices were baked onto a glass slide, and heat-stained with 0.5% aqueous toluidine blue. For myelinated axon count, images were captured using a Nikon Eclipse E600 microscope with 40× and 100× objectives. The total number of myelinated axons in each nerve was counted automatically in ImageJ (v1.52p) and manually verified as previously described (Bogdanik *et al*, 2013). It was determined that there were no sex differences in phenotype, and therefore the data was presented as mixed sex, with similar numbers of males and females. Nerve images were generated by stitching together images encompassing the whole field using the ImageJ (v1.52p) Stitching Grid/Pairwise plugin (Preibisch *et al*, 2009).

### Denervation

Medial gastrocnemius (MG) muscle was dissected and fixed in freshly prepared 2% paraformaldehyde (Electron Microscopy Sciences) in phosphate-buffered saline (PBS) overnight. Samples were then incubated in blocking solution [2.5% bovine serum albumin (Sigma-Aldrich) and 1% Triton-X 100 (Sigma-Aldrich) in PBS] for 1h, before being gently teased apart and pressed between two glass slides using a binder clip, for 15 min at 4◻°C. Samples were subsequently returned to the blocking solution, permeabilized overnight at 4◻°C, and incubated with primary antibodies [1:500 mouse monoclonal IgG1 anti-neurofilament 2H3 and 1:250 anti-SV2 (both obtained from the Developmental Studies Hybridoma Bank) in blocking buffer] overnight at 4◻°C on a slow shaker. Samples were then washed at least 4 times for 15 min in PBS and incubated overnight at 4◻°C on a slow shaker with Alexa-Fluor 488 goat anti-mouse IgG1 and α-bungarotoxin conjugated with Alexa-Fluor 594 (1:500 and 1:1,000 dilutions, respectively. Both from Invitrogen, Carlsbad, CA) to stain for acetylcholine receptor (AChR). After incubation, samples were rinsed three times and washed at least four times for 15 min each, mounted in 80% glycerol, and imaged using an SP5 Leica confocal microscope. Occupancy of 50 or greater randomly selected neuromuscular junctions (NMJs) was scored blinded to genotype on a Nikon E600 fluorescence microscope. Full occupancy of NMJ was defined as when presynaptic nerve staining fully overlaid with AChR, partial occupancy as when positive for AChR but only partially stained for presynaptic nerve, and denervation as when positive for AChR but negative for presynaptic nerve staining. It was determined that there were no sex differences in phenotype and therefore the data was presented as mixed sex, with similar numbers of males and females.

### KLHDC10/Pirh2 Co-Immunoprecipitation

Four million HeLa cells of indicated genotypes were plated in 10-cm dishes and transfected the following day with two plasmids: one expressing an Ala-tail-binding E3 ligase (3xFLAG-KLHDC10 or -Pirh2) and the other, a ribosome stalling or control reporter (GFP-NS or GFP, respectively) in a 1:1 mass ratio. The next day, cells were treated with 10 μM MG-132 for 4h and then washed in ice-cold PBS before lysis. Cell lysis was conducted in ice-cold β-Octyl glucoside lysis buffer (20 mM Tris-HCl pH 7.5, 100 mM KOAc, 5 mM MgCl_2_, 1% β-Octyl glucoside, 1 mM DTT) supplemented with EDTA-free protease inhibitor tablet (Roche), for 30 min at 4°C, followed by lysate clarification by centrifugation (12,000 x *g*, 5 min, 4 ºC). GFP was immunoprecipitated from lysates by adding 10 μl GFP-Trap magnetic agarose beads (ChromoTek) and incubating for 3h at 4 ºC. Beads were washed 3x in lysis buffer without DTT before elution by boiling the beads in 2X Laemmli buffer. Both whole-cell lysates (WCL) and immunoprecipitation eluates were analyzed by Western blot (WB).

### Cell lines

HeLa cells deleted for *LTN1, NEMF*, or both, have been previously described (Thrun *et al*., 2021). HeLa cells were maintained in DMEM (Gibco) supplemented with 10% FBS (Gibco), penicillin and streptomycin. Antibiotics were omitted from the culture media during experiments. Cells were transfected with polyethylenimine STAR (PEI, Tocris) in a 1:4 DNA:PEI mass ratio, or alternatively, with Lipofectamine 3000 (Invitrogen) following the manufacturer’s instructions. When indicated, cells were treated with 10 µM MG-132 in DMSO for 4h.

For all NEMF reconstitution experiments, we employed cells stably overexpressing NEMF, which were produced by lentiviral infection (see below). These cells were maintained as above, but with the addition of puromycin selection (1 µg/ml). Puromycin was omitted during experiments. To detect aggregate formation (Fig. 4E), we used a transient rescue approach that led to physiological levels of NEMF re-expression.

### Lentivirus production and infection

Lentiviral particles were produced in HEK-293T in a biosafety level 2 laboratory. Cells were maintained as described above. Four million HEK-293T cells in a 10-cm plate were transfected with 15 µg of the plasmids pWPI-NEMF-5XFLAG-IRES-puro (encoding WT or D96A mutant NEMF), pCMV-Δ8.91 and pMD2.G (3:3:1 mass ratio) using Lipofectamine 3000, and the medium was replaced 8h later. Forty-eight hours post-transfection, the supernatant was collected, centrifuged (200 x *g*, 5 min) and filtered through 0.22 µm pore-size membranes. This supernatant was used to infect HeLa *N-KO* or *LN-KO* twice. Cells were then selected with puromycin (2 µg/ml) for a week before expansion.

### Cell lysis

Cells were washed once with ice-cold PBS followed by lysis in NP-40 buffer (20 mM Tris-HCl pH 7.5, 150 mM NaCl, 5 mM EDTA, 0.5% NP-40, 1 mM DTT), supplemented with EDTA-free protease inhibitor tablet (Roche) (Fig. 4F), or in NP-40-Glycerol buffer (50 mM Tris-HCl pH 7.5, 100 mM NaCl, 10 mM MgCl_2_, 0.1% NP-40, 5% glycerol, 1 mM DTT), supplemented with EDTA-free protease inhibitor tablet (Roche) (other figures), for 5 min at 4°C. Insoluble cellular debris was discarded by centrifugation at 12,000 x *g* for 10 min at 4°C and the supernatants collected. Laemmli buffer was added to lysates, samples were boiled for 5 min at 95°C and analyzed by immunoblotting as described below.

### Western blotting

Protein samples were resolved on precast SurePAGE Bis-Tris 4-20% gradient gels (GenScript) or 16% Tris-Glycine gels (Invitrogen) (Fig. 4F) following the supplier’s guidelines. Proteins were transferred to PVDF membranes (0.2 µm or 0.45 µm pore size) using the Turbo-BLOT semi-dry transfer system with Mixed Mw program settings (Bio-Rad) or employing the Mini-Blot transfer system (Invitrogen) using SDS-Towbin’s transfer buffer (20% Methanol, 25 mM Tris-HCl, 192 mM Glycine, 0.1% SDS) for 1.5h at 35V (Fig. 4F). For protein detection, membranes were blocked in 5% non-fat milk for 1h at room temperature, and subsequently incubated with the indicated antibodies (2h at room temperature or overnight at 4 ºC) and HRP-labeled secondary antibodies. Protein bands were developed in an ImageQuant 800 device (Cytiva) using the SuperSignal™ West Pico PLUS Chemiluminescent Substrate or the SuperSignal™ West Femto Maximum Sensitivity Substrate (Thermo Fisher).

### Plasmids

The following plasmids were used in this study:

| Plasmid name | Source |
| --- | --- |
| pCMV-5FBG-stop | Gift (Saito *et al.*, 2013) |
| pCMV-5FBG-Nonstop | Gift (Saito *et al.*, 2013) |
| pEGFP-stop | ClonTech (discontinued) |
| pEGFP-Nonstop | (Thrun *et al.*, 2021) |
| pcDNA5-3XFLAG-KLHDC10 | (Thrun *et al.*, 2021) |
| pcDNA5-3XFLAG-Pirh2 | (Thrun *et al.*, 2021) |
| pWPI-IRES-Puro | (Magg *et al.*, 2024) |
| pCMV-Δ8.91 | (Magg *et al.*, 2024) |
| pMD2.G | (Magg *et al.*, 2024) |
| pWPI-NEMF-5XFLAG-IRES-Puro | This study |
| pWPI-NEMF-D96A-5XFLAG-IRES-Puro | This study |
| pcDNA5-NEMF-FLAG | This study |
| pcDNA5-NEMF-D96A-FLAG | This study |
| pCMV-6His-4FBG(KR)-Nonstop | This study |

### Antibodies

The following antibodies and dilutions were used in this study:

Antibodies The following antibodies and dilutions were used in this study:
| **Antibody** | **Source (Catalog number)** | **Dilution** |
| --- | --- | --- |
| anti-GFP | Roche (Cat. No. 11814460001) | 1:1,000 |
| anti-GFP | MBL (Cat. No. 598) | 1:1,000 |
| anti-FLAG M2 | Sigma-Aldrich (Cat. No. F1804) | 1:1,000 |
| anti-GAPDH | CST (Cat. No. 2118) | 1:1,000 |
| anti-RPL4 | ProteinTech (Cat. No. 11302-1-AP) | 1:2,000 |
| anti-NEMF | ProteinTech (Cat. No. 11840-1-AP) | 1:1,000 |
| anti-Listerin | Thermo Fisher (Cat. No. PA5-42315) | 1:1,000 |
| Anti-6xHis | Thermo Fisher (Cat. No. MA1-21315) | 1:1000 |

### Statistical analyses

Data are presented as mean ± SD unless otherwise indicated. The statistical tests applied are described in each figure legend, and whenever possible, exact p-values are provided. In vivo experiments were conducted in a blind manner with respect to mouse genotype. It was determined that there were no sex differences in phenotype (except for body weight) and therefore the data was presented as mixed sex, with similar numbers of males and females. The sample size for each experiment is indicated in the respective figure legend.

## Results

### NEMF-D96A mutation impairs Ala-tailing activity

Previous studies have pinpointed the NEMF D96 residue as essential for NEMF-mediated Ala-tailing (Udagawa *et al*., 2021; Thrun *et al*., 2021). To verify and further extend these observations, we first evaluated the effect of the NEMF D96A mutation on the Ala-tailing of an RQC-dependent reporter according to its mobility shift in SDS-PAGE. This reporter consists of a 5xFLAG tag fused to the N-terminus of the β-globin gene (5FBG) lacking a stop codon (hereafter, 5FBG-NS) (Saito *et al*., 2013) (Fig. 1A, schematic). Translation through the β-globin gene into the polyA tail causes ribosome stalling and subsequent RQC-mediated degradation when expressed in wild-type (*wt*) HeLa cells (Fig. 1A, compare lanes 1 vs 5). That the reporter readily becomes Ala-tailed is indicated by the appearance of a smear migrating immediately above the full-length product in SDS-PAGE (Fig. 1A, lower panel, compare lanes 1 vs 5) that is strictly dependent on NEMF, as it disappears in NEMF-KO HeLa cells (hereafter, ‘HeLa *N-KO*’) (Fig. 1A, lane 6). Moreover, re-expression of NEMF in HeLa *N-KO* cells, but not the D96A mutant, restored Ala-tailing, thus confirming that the mutation indeed impairs NEMF’s enzymatic activity (Fig. 1A, lanes 7 and 8) and (Udagawa *et al*., 2021). Importantly, the NEMF D96A mutant was expressed at normal levels (Fig. 1A, lanes 7 and 8).

**Figure 1.**
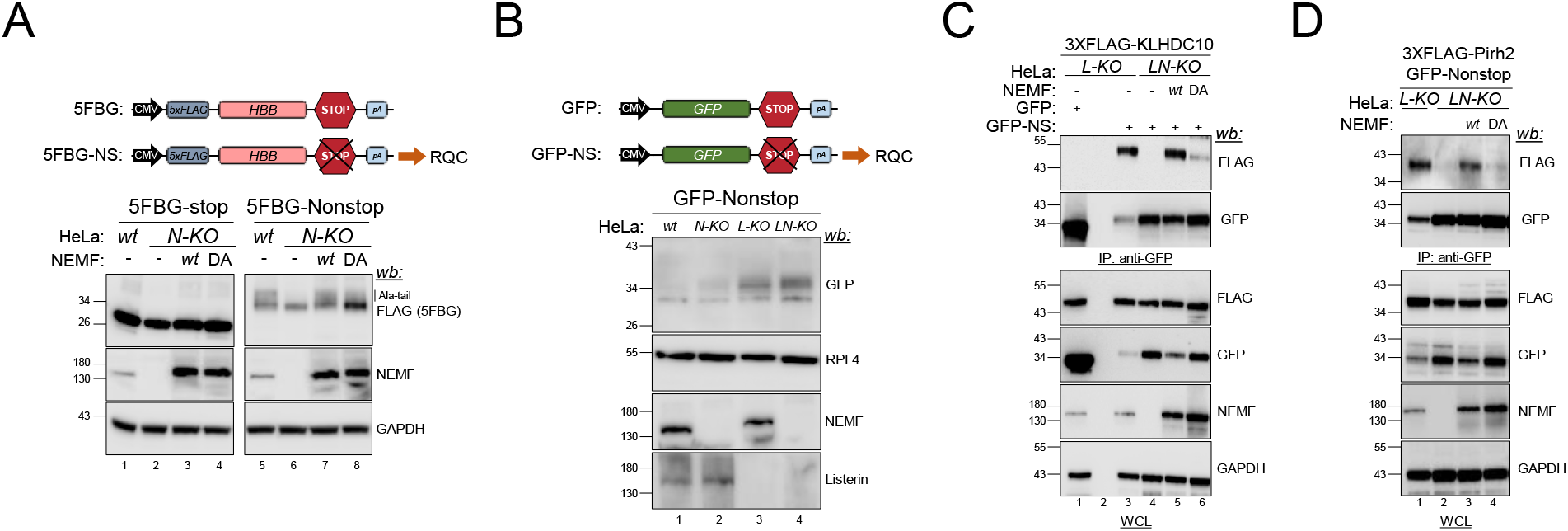
The *Nemf*-D96A mutation impairs Ala-tailing. A. Schematic of the RQC-dependent reporter 5FBG-NS. HeLa cells of indicated genotypes (*wt, N-KO* or *N-KO* reconstituted with *wt* or D96A NEMF) were transfected with the 5FBG-NS reporter. Immunoblots: anti-FLAG and anti-NEMF to monitor reporter and NEMF levels, respectively, and anti-GAPDH as loading control. A representative experiment is shown (n = 3). B. Schematic of the RQC-dependent reporter GFP-NS. HeLa cells of indicated genotypes (*wt, N-KO, L-KO* or *LN-KO*) were transfected with the GFP-NS reporter. Immunoblots: anti-GFP, anti-NEMF and anti-Listerin to monitor reporter, NEMF and Listerin levels, respectively and anti-RPL4 as loading control. A representative experiment is shown (n = 3). C. The interaction of GFP and GFP-NS with 3XFLAG-tagged KLHDC10 in HeLa cells of indicated genotypes (*L-KO, LN-KO* or *LN-KO* reconstituted with *wt* or D96A NEMF) was analyzed by GFP-Trap. Immunoblots: anti-GFP, anti-FLAG and anti-NEMF to monitor reporter, KLHDC10 and NEMF levels, respectively, and anti-GAPDH as loading control. A representative experiment is shown (n = 3). D. The interaction of GFP and GFP-NS with 3XFLAG-tagged Pirh2 in HeLa cells of indicated genotypes (*L-KO, LN-KO* or *LN-KO* reconstituted with *wt* or D96A NEMF) was analyzed by GFP-Trap. Immunoblots: anti-GFP, anti-FLAG and anti-NEMF to monitor reporter, Pirh2 and NEMF levels, respectively, and anti-GAPDH as loading control. A representative experiment is shown (n = 3).

Next, as an orthogonal approach to further assess the effect of the NEMF D96A mutation on Ala-tailing activity, we co-opted the ability of the E3 ligases KLHDC10 and Pirh2 to selectively bind Ala-tail degrons (Thrun *et al*., 2021). The assay consisted of monitoring the Ala-tailing-dependent co-immunoprecipitation of KLHDC10 with an RQC reporter, GFP lacking stop codons (hereafter, ‘GFP-NS’; Fig. 1B, schematic). As RQC reporters are targeted for proteasomal degradation by both NEMF and Listerin [Fig. 1B, and (Thrun *et al*., 2021)], to increase GFP-NS steady-state levels and facilitate detecting the interaction, we used proteasome-inhibited and *LTN1*-KO HeLa cells (hereafter, ‘HeLa *L-KO*’). In HeLa *L-KO*, KLHDC10 co-immunoprecipitated with GFP-NS, but not with GFP, despite the substantially reduced expression of GFP-NS due to ribosome stalling (Bengtson & Joazeiro, 2010) (Fig. 1C, lanes 1 and 3). Knocking out *NEMF* in the *L-KO* background (hereafter, ‘HeLa *LN-KO*’) severely diminished KLHDC10 binding, an effect that could be rescued by *wt* NEMF reconstitution, but not by the D96A mutant (Fig. 1C, lanes 3 to 6). Comparable results were obtained when Ala-tailing was assessed via Pirh2 binding under a similar experimental setting (Fig. 1D). Together, the above reporter mobility-shift and KLHDC10/Pirh2 co-immunoprecipitation data confirm that the NEMF D96A mutation led to defective Ala-tailing.

### Nemf D96A homozygous mice show a mild motor phenotype

To determine the physiological contribution of NEMF-mediated Ala-tailing to translational homeostasis, we generated homozygous *Nemf*-D96A knock-in mice using CRISPR–Cas9 technology (nucleotide: A289→C). Homozygous mice bearing the D96A mutation (C57BL/6J^*Nemf*-D96A/D96A^, hereafter *Nemf* ^D96A/D96A^) were viable and obtained at the expected mendelian ratio, grew normally, as measured by weight gain, and had a similar lifespan compared to that of *wt* littermates (Fig. 2A and B).

**Figure 2.**
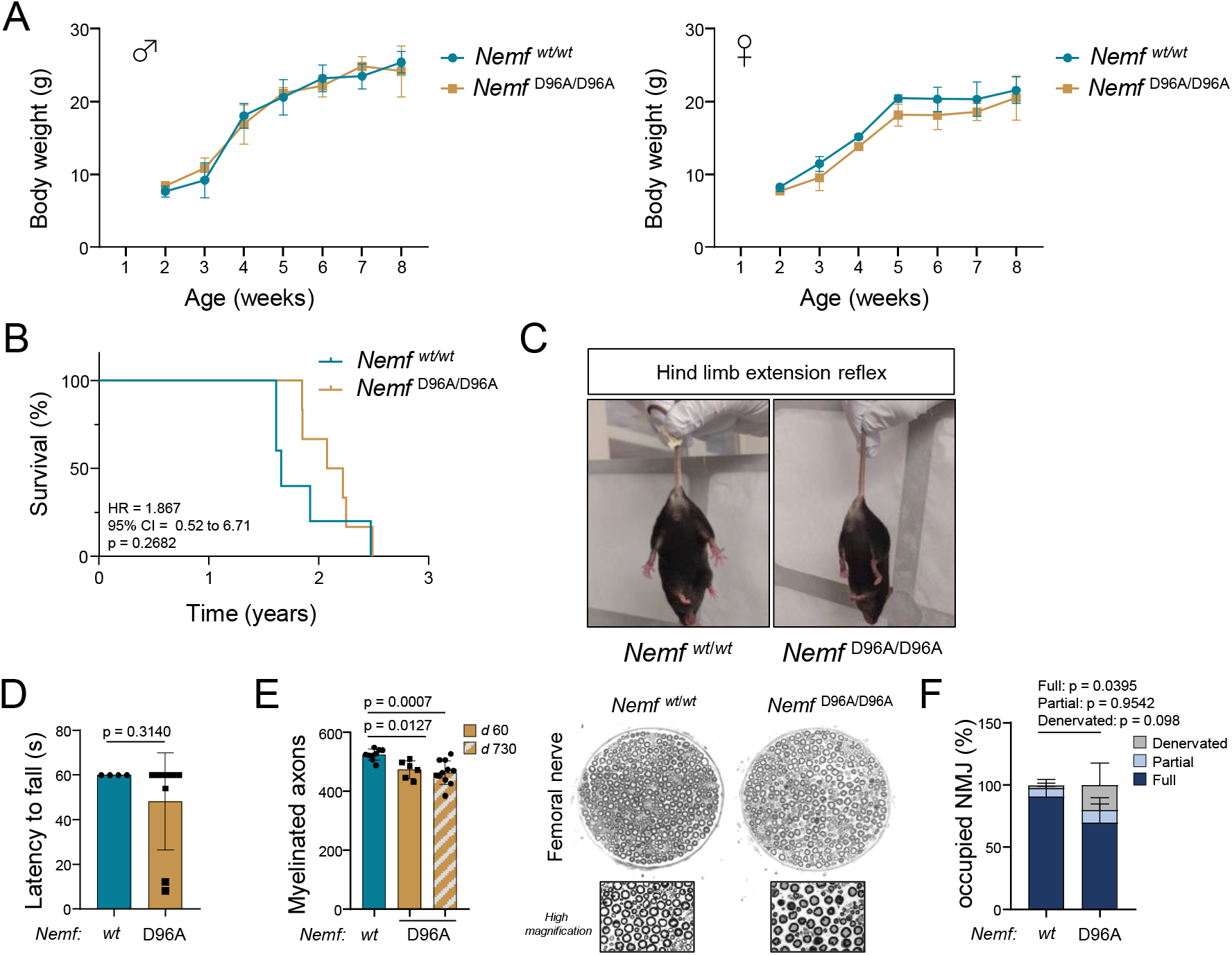
*Nemf*-D96A homozygous mice have a mild motor phenotype. A. Body weight gain (in g) over time (in weeks) in male *Nemf* ^D96A/D96A^ (left) or female *Nemf* ^D96A/D96A^ (right) compared to sex-matched *wt* counterparts (n = 3-6 mice). B. Survival curve of *Nemf* ^D96A/D96A^ or *wt* littermates (n = 5 mice for *wt*, 6 mice for *Nemf* ^D96A/D96A^). p-values were obtained by log-rank test. HR: Hazard ratio; CI: Confidence interval. *C. Nemf* ^D96A/D96A^ (right image) show a defective hind limb extension reflex when suspended from the tail. D. Latency to fall (in s) in the wire hang test. p-values were obtained by unpaired Student’s *t* test (n = 4 mice for *wt*, 9 mice for *Nemf* ^D96A/D96A^). E. Myelinated axon count in the femoral nerve motor branch in adult (*d* 90, i.e. 3 months-old; n = 6 mice) or aged (*d* 730, i.e. 2 years-old; n = 11 mice) *Nemf* ^D96A/D96A^ mice compared to adult *wt* mice (n = 9 mice). Representative cross-sections of femoral nerve motor branches and their magnifications are shown. p-values were obtained by one-way ANOVA with Dunnett’s post-hoc correction. F. Quantification of fully innervated, partially innervated or denervated NMJs based on presynaptic and postsynaptic staining overlap. *wt* and *Nemf* ^D96A/D96A^ (n◻= 5, and 3 mice, respectively) were analyzed. p-values were obtained by two-way ANOVA with Tukey’s post-hoc correction.

We had previously identified NEMF mutations from N-ethyl-N-nitrosourea (ENU) mutagenesis that cause progressive motor neuron degeneration and motor impairment in mice (Martin *et al*., 2020). Therefore, we analyzed motor phenotypes in this new *Nemf* ^D96A/D96A^ mouse model. Adult homozygous mice [*ca*. post-natal day (*d*) 90] exhibited impaired hind limb extension reflex when suspended from the tail (Fig. 2C), but similar muscle strength to that of *wt* mice, as measured by the latency to fall in the wire hanging test (Fig. 2D). Consistent with this mild defect, histological examination of the femoral nerve showed only a minor reduction in myelinated motor axons which did not progress further with age until *d* 730 (Fig. 2E). In contrast to the more severe phenotypes observed in our previously-reported *Nemf* ^R86S/R86S^ ENU mutant mouse (Martin *et al*., 2020), the small decrease in myelinated axon count in *Nemf* ^D96A/D96A^ mice was accompanied by a similarly mild loss of the integrity of neuromuscular junctions (NMJs) (Fig. 2F). Overall, these data indicate that the NEMF-D96A mutation in mice causes a mild motor phenotype.

### Loss of Ala-tailing activity aggravates the lister phenotype

We sought to obtain evidence for a critical physiological role of Ala-tailing by leveraging the mild phenotype caused by the *Nemf-*D96A mutation to examine genetic interactions with *Ltn1*. If failure of RQC plays a crucial role in neurodegeneration of *lister* mice, and if Ala-tailing is relevant for translational surveillance, the model predicts that inactivation of Ala-tailing, by affecting both proteolytic mechanisms of RQC, namely RQC-L and RQC-C (Thrun *et al*., 2021), would lead to a more severe phenotype compared to the *Ltn1* mutation alone. To test this hypothesis, we generated mice with a combination of the *lister* mutation (Chu *et al*., 2009) and the *Nemf*-D96A allele.

First, we intercrossed *Ltn1* ^*lister*/*wt*^;*Nemf* ^D96A/*wt*^ mice and analyzed the genotype distribution of the resulting offspring. Out of 197 born and genotyped mice, none were double-homozygous *Ltn1* ^*lister*/*lister*^;*Nemf* ^D96A/D96A^ although *ca*. 12 were expected by Mendelian distribution (Figure 3A, p-value between distributions = 0.0008). To increase the potential yield of double-homozygous animals, we next mated *Ltn1* ^*lister*/*wt*^;*Nemf* ^D96A/*wt*^ to *Ltn1* ^*lister*/*wt*^;*Nemf* ^D96A/D96A^ mice. Again, out of 46 descendant offspring analyzed, none had the *Ltn1* ^*lister*/*lister*^;*Nemf* ^D96A/D96A^ genotype combination, although ∼6 were expected (Figure 3B, p-value between distributions = 0.0578). Finally, we intercrossed *Ltn1* ^*lister*/*wt*^;*Nemf* ^D96A/D96A^ mice littermates to further enforce inheritance of the *Nemf*-D96A allele. In this experiment, we closely monitored the offspring from birth, and observed that only perinatally deceased mice (*d* 1) had the *Ltn1* ^*lister*/*lister*^;*Nemf* ^D96A/D96A^ genotype. Although the double-mutant homozygous animals were observed at half the expected frequency, this difference was not statistically significant. Therefore, although one cannot determine if the double mutation affects embryonic development, these animals clearly cannot survive post-natally (Figure 3C, p-value between distributions of all animals = 0.0799; p-value excluding dead animals < 0.0001).

**Figure 3.**
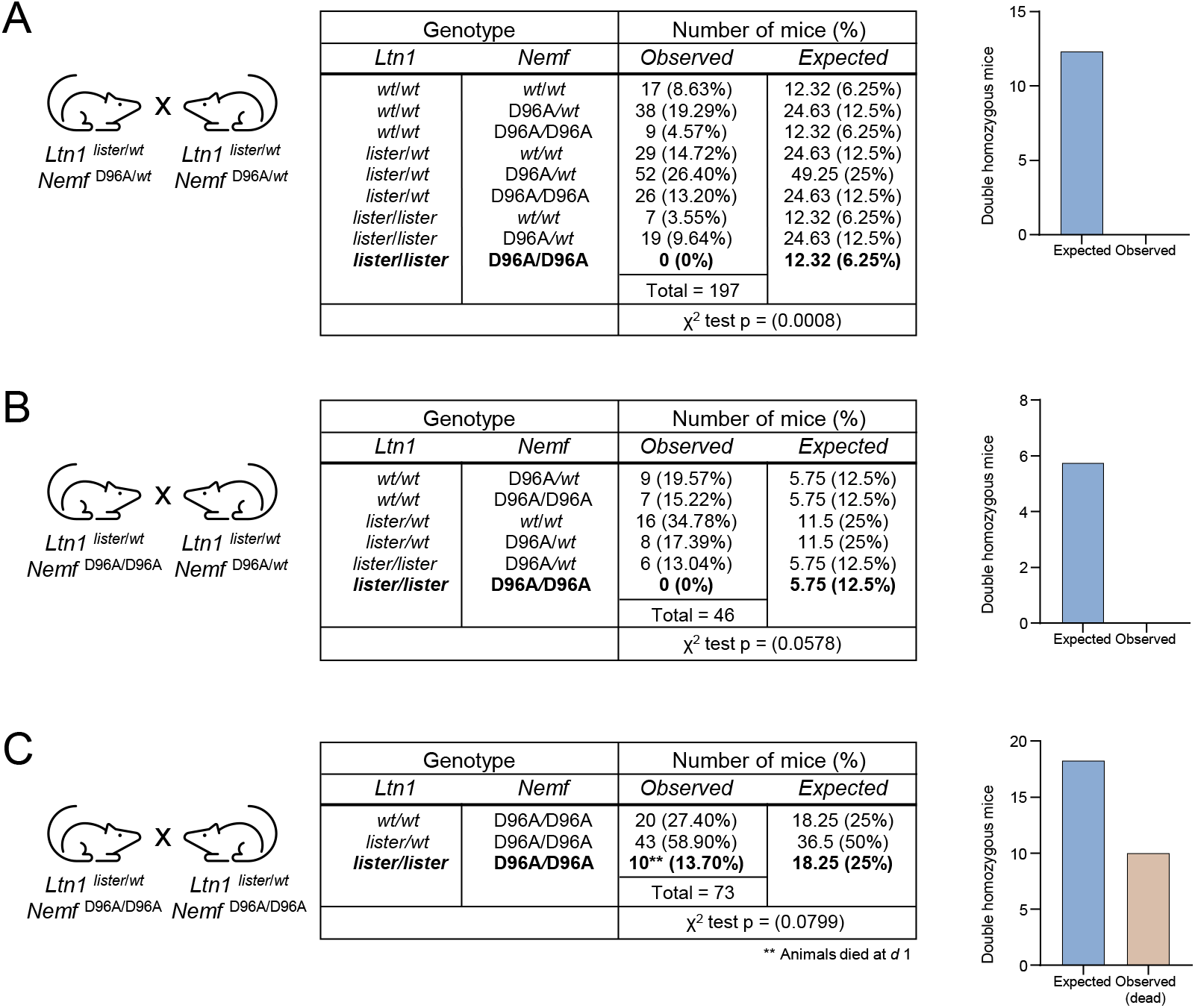
Genetic inactivation of Ala-tailing enhances the severity of the *lister* phenotype. A. Resultant offspring and genotypes from *Ltn1* ^*lister*/*wt*^;*Nemf* ^D96A/*wt*^ x *Ltn1* ^*lister*/*wt*^;*Nemf* ^D96A/*wt*^ crosses. None double homozygous mice (in bold) were obtained, despite expecting 6.25% of the descendants (∼12 animals). p-values between observed and expected distributions were obtained by χ^2^-test. B. Resultant offspring and genotypes from *Ltn1* ^*lister*/*wt*^;*Nemf* ^D96A/D96A^ x *Ltn1* ^*lister*/*wt*^;*Nemf* ^D96A/*wt*^ crosses. None double homozygous mice (in bold) were obtained, despite expecting 12.5% of the descendants (∼6 animals). p-values between observed and expected distributions were obtained by χ^2^-test. C. Resultant offspring and genotypes from *Ltn1* ^*lister*/*wt*^;*Nemf* ^D96A/D96A^ x *Ltn1* ^*lister*/*wt*^;*Nemf* ^D96A/D96A^ crosses. Ten double homozygous mice that died at *d* 1 (in bold) were obtained, despite expecting 25% of the descendants (∼18 animals). p-values between observed and expected distributions were obtained by X^2^-test.

Overall, these results indicate a strong synthetic defect in mice caused by simultaneous impairment of Listerin and NEMF’s Ala-tailing functions. This provides robust evidence supporting the relevance of Ala-tailing-mediated nascent chain degradation for translational homeostasis *in vivo*.

### Genetic reduction of Ala-tailing partially rescues lister phenotypes

During the course of the above genetic interaction experiments, we noticed a remarkable phenotypic improvement in the *lister* phenotype when mice homozygous for the *Ltn1* mutation also inherited a single copy of the *Nemf*-D96A allele. Namely, *Nemf*-D96A heterozygosity in *lister* mice partially recovered animal growth (Fig. 4A), and most notably, expanded lifespan ∼6-fold, from a median of 53 days to 313 days (Fig. 4B). *lister* mice display age-progressive motor neuron loss, which manifests, among other behavioral alterations, as a defective hind limb extension reflex. However, *lister* animals with a heterozygous *Nemf*-D96A mutation did not display any impairment in this response (Fig. 4C). Additionally, *lister* mice performed progressively worse in the wire hang test (Fig. 4D), whereas *Ltn1* ^*lister*/*lister*^;*Nemf* ^D96A/*wt*^ mice did not show any phenotypic trait in this test at any timepoint analyzed, strongly suggesting that these animals maintain normal motor function (Fig. 4D). Taken together, these findings suggest that Ala-tailing is a key mechanism driving neurodegeneration in *lister* mice.

**Figure 4.**
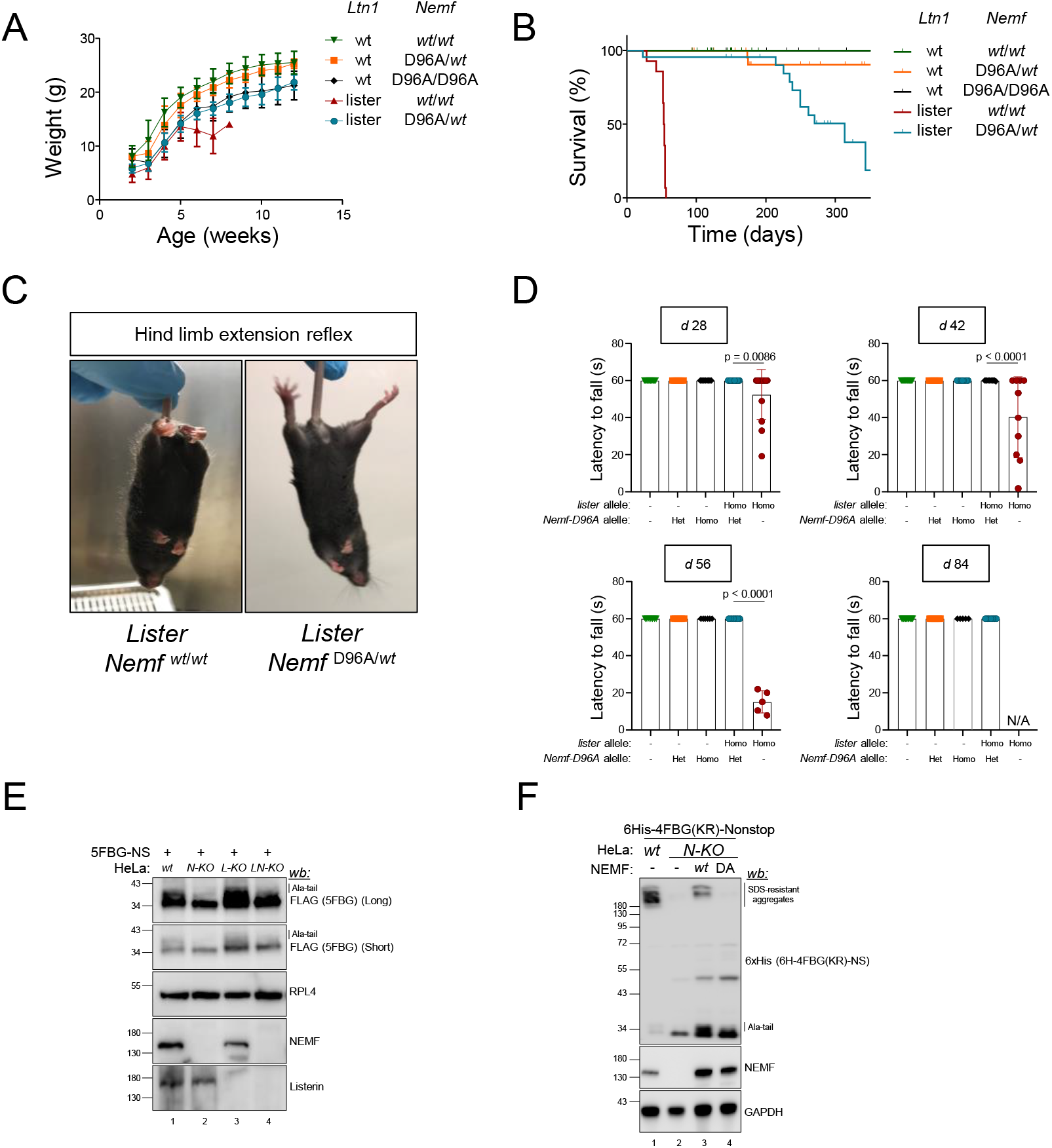
Ala-tailing contributes to toxicity caused by Listerin mutation. A. Body weight gain (in g) over time (in weeks) of mice with the indicated genotypes. *Nemf*-D96A heterozygous *lister* animals display a partially recovered phenotype (n = 7 – 11 mice per group). B. Survival curves of mice with the indicated genotypes. *Nemf*-D96A heterozygous *lister* animals display an extended lifespan (n = 14 – 29 mice per group). *C. Lister* animals (left image) show a defective hind limb extension reflex when suspended from the tail that is rescued by heterozygous *Nemf*-D96A (right image). D. Latency to fall (in s) in the wire hang test in *d* 28, 42, 56 and 84. Missing points are due to deceased animals. p-values were obtained by one-way ANOVA with Tukey’s post-hoc correction (n = 5 – 17 mice per group and timepoint). E. HeLa cells of indicated genotypes (*wt, N-KO, L-KO* or *LN-KO*) were transfected with the 5FBG-NS reporter. Immunoblots: anti-FLAG, anti-Listerin and anti-NEMF to monitor reporter, Listerin and NEMF levels, respectively, and anti-RPL4 as loading control. A representative experiment is shown (n = 3). F. HeLa cells of indicated genotypes (*wt, N-KO* or *N-KO* reconstituted with *wt* or D96A NEMF) were transfected with the 6His-4FBG(KR)-NS reporter. Immunoblots: anti-FLAG and anti-NEMF to monitor reporter and NEMF levels, respectively, and anti-GAPDH as loading control. A representative experiment is shown (n = 3).

### Accumulation of Ala-tailed proteins contributes to cytotoxicity in lister mice

Next, we set out to investigate why partially reducing Ala-tailing in *lister* mice was beneficial. Previous work had shown that in yeast deleted for the Listerin ortholog, Ltn1, increased C-terminal-tailing by Rqc2 (the NEMF ortholog) caused protein aggregation and cytotoxicity (Yonashiro *et al*., 2016; Choe *et al*., 2016; Sitron *et al*, 2020). Accordingly, knockdown of NEMF increased both the solubility of an overexpressed nonstop reporter and cellular viability in HeLa cells (Udagawa *et al*., 2021). We therefore investigated if Ala-tailing plays a role in the aggregation of RQC substrates in mammalian cells. Expression of the RQC reporter 5FBG-NS in *wt* HeLa cells was accompanied by its modification with an Ala-tailing smear, which was absent in *N-KO* cells (Figs. 1A and 4E). Consistent with yeast data, the levels of Ala-tailed reporter were increased in Listerin-deficient (*L-KO*) cells compared to *wt* cells (Fig. 4E). However, under these conditions we failed to observe reporter aggregation in SDS-PAGE. Rationalizing that detecting reporter aggregation might require its further accumulation, we generated a variant reporter in which β-globin Lys residues were replaced with Arg, with the expectation to simultaneously interfere with the action of the RQC E3 ligases, Listerin, Pirh2 and KLHDC10. Strikingly, this reporter formed conspicuous SDS-resistant aggregates in *wt* cells (Fig. 4F). Moreover, this aggregation was NEMF-dependent, as it could not be observed in *N-KO* cells. Finally, re-expression of NEMF *wt* but not the D96A mutant, restored aggregate formation, indicating this is an Ala-tail-dependent mechanism (Fig. 4F). Collectively, these observations provide a plausible explanation for the phenotypic rescue of lister mice by reducing Ala-tailing activity, suggesting that accumulation and aggregation of Ala-tailed proteins majorly contributes to neurodegeneration caused by Listerin mutation.

## Discussion

In this study, we developed a new mouse model with NEMF’s Ala-tailing activity selectively impaired. The results of mouse genetic and biochemical analyses provide evidence for Ala-tailing’s critical role in translational surveillance in both health and disease. Moreover, we have gained insights into the long-standing question of how *Ltn1* mutation causes neurodegeneration. The evidence implicates the accumulation and aggregation of Ala-tailed proteins that evade elimination as a major contributor to the pathomechanism.

*Nemf*-D96A homozygous mice were viable and overall healthy. One possibility is that the mutant protein retains residual Ala-tailing activity. Alternatively, Ala-tailing may not be essential for developmental and physiological translation surveillance due to Listerin’s predominant role in RQC-mediated proteolytic targeting. Since the D96A mutation, as the equivalent mutation in NEMF orthologs, does not impact ribosome or Listerin binding (Shen *et al*., 2015; Yonashiro *et al*., 2016; Choe *et al*., 2016; Lytvynenko *et al*., 2019; Udagawa *et al*., 2021; Takada *et al*., 2021), the association of Listerin to obstructed 60S subunits can still be stabilized by NEMF D96A, allowing Listerin-mediated ubiquitination of aberrant nascent chains. Consistent with this model, although the D96A mutation had little effect on its own, it enhanced the phenotype caused by the *Ltn1* mutation—notably, the early lethality of *Ltn1* ^*lister*/*lister*^;*Nemf* ^D96A/D96A^ mice is reminiscent of the phenotype caused by *Nemf* knockout (Martin *et al*., 2020), which fully inactivates NEMF function while also interfering with Listerin function.

Our data shows that Ala-tailing was critical for the post-natal survival of *lister* mice up to early adulthood (∼60 days), supporting a compensatory role for Ala-tail-dependent proteolytic mechanisms during this period. However, the question remains as to why Ala-tailing does not provide long-term compensation in *lister* mice. Notably, *lister* mice with partially reduced Ala-tailing (*Ltn1* ^*lister*/*lister*^;*Nemf* ^D96A/*wt*^) are healthier and live longer than *lister* mice with two *wt Nemf* alleles. This finding suggests that, at some point post-natally, Ala-tailed proteins become toxic due to an inability of the cellular machinery to handle them. One plausible explanation is that the capacity of the E3 ligases in RQC-C is exceeded, leading to the accumulation of Ala-tailed proteins. Consistently, in a previous study, overexpression of GFP hard-coded with C-terminal Ala residues led to its aggregation, and cytotoxicity in HeLa cells (Udagawa *et al*., 2021). Thus, a decrease in Ala-tail-mediated protein aggregation represents a plausible mechanism to explain the improved outcomes of *Ltn1* ^*lister*/*lister*^;*Nemf* ^D96A/*wt*^ mice, as well as why Ala-tailing eventually fails to compensate for Listerin’s loss-of-function. Overall, the results reveal a striking duality of Ala-tailing, being both neuroprotective and neurotoxic.

In humans, expansion of polyalanine has been identified in nine genes, all of which cause aggregation of the mutant protein and congenital diseases, including neurological disorders (Hughes & Thomas, 2013). Although these poly-alanine tracts may form aggregates that are not amyloid (Polling *et al*, 2015), unlike what we observe for Ala-tails, their similar poly-alanine composition warrants an investigation on the extent to which pathogenic mechanisms overlap, between Ala-tail toxicity in *lister* mice and these inherited poly-alanine diseases.

A fascinating question that arises is why Listerin dysfunction and the resulting aggregation of Ala-tailed proteins are particularly detrimental to neuronal cells. One possibility is that these aggregates, similarly to those occurring in major neurodegenerative diseases, sequester proteins essential for neuronal function (Wilson *et al*, 2023). In yeast, Rqc2-dependent aggregates sequester chaperones, perturbing global proteostasis and cell viability (Choe *et al*., 2016; Yonashiro *et al*., 2016; Sitron *et al*., 2020). Alternatively, given their complex and polarized morphology, neurons could be hypersensitive to Ala-tail-dependent and other types of aggregates. Thus, elucidating the protein composition and consequences of Ala-tail-dependent aggregates is a critical area for future research.

In conclusion, our results provide evidence, for the first time, for the crucial role of Ala-tailing in translational surveillance and global proteostasis in a mammalian organism. Using *lister* mice, which experience translational stress, we demonstrate that the formation of Ala-tails must be finely tuned to prevent toxic protein accumulation and aggregation. Given the emerging evidence showing the prevalence of impaired translation landscapes in various diseases, we propose that deciphering the pathophysiological roles of RQC in different contexts will provide new insights into the molecular basis of other pathological conditions.

## Declaration of interest

The authors declare no competing interests.

## Acknowledgments

We thank the ZMBH FACS Facility, and the Joazeiro laboratory for discussions. We also thank Dr. Alessia Ruggieri and Prof. Shin-ichi Hoshino for constructs. Work in the Joazeiro laboratory is supported in part by the Deutsche Forschungsgemeinschaft (DFG, German Research Foundation) Project 201348542 (SFB 1036) and by the National Institute of Neurological Disorders and Stroke (NINDS) of the NIH (R01 NS102414) to GAC and CAPJ.

## Notes

### Competing Interest Statement

The authors have declared no competing interest.

